# An approachable, flexible, and practical machine learning workshop for biologists

**DOI:** 10.1101/2022.02.03.479008

**Authors:** Chris S Magnano, Fangzhou Mu, Rosemary S Russ, Milica Cvetkovic, Debora Treu, Anthony Gitter

## Abstract

The increasing prevalence and importance of machine learning in biological research has created a need for machine learning training resources tailored towards biological researchers. However, existing resources are often inaccessible, infeasible, or inappropriate for biologists because they require significant computational and mathematical knowledge, demand an unrealistic time-investment, or teach skills primarily for computational researchers. We created the Machine Learning for Biologists (ML4Bio) workshop, a short, intensive workshop that empowers biological researchers to comprehend machine learning applications and pursue machine learning collaborations in their own research. The ML4Bio workshop focuses on classification and was designed around 3 principles: (a) focusing on preparedness over fluency or expertise, (b) necessitating minimal coding and mathematical background, and (c) requiring low time investment. It incorporates active learning methods and custom open source software that allows participants to explore machine learning workflows. After multiple sessions to improve workshop design, we performed a study on 3 workshop sessions. Despite some confusion around identifying subtle methodological flaws in machine learning workflows, participants generally reported that the workshop met their goals, provided them with valuable skills and knowledge, and greatly increased their beliefs that they could engage in research that uses machine learning. ML4Bio is an educational tool for biological researchers, and its creation and evaluation provides valuable insight into tailoring educational resources for active researchers in different domains. The workshop materials are available from https://carpentries-incubator.github.io/ml4bio-workshop/ and the ml4bio software is available from https://github.com/gitter-lab/ml4bio.

## 1 Introduction

Machine learning (ML) is a powerful tool for analyzing biological data and is increasingly popular in biological research. Biological publications using ML have increased exponentially over the past decades^1^. In 2017 almost 90% of 704 NSF principal investigators reported that they “are currently or will soon be analyzing large datasets.”^2^ However, the most commonly reported unmet needs were training-based. As of 2017, only about a quarter of life-sciences training programs taught necessary skills for data stewardship^3^. In the United States, the National Science Foundation and National Institutes of Health have recognized the need for training at the intersection of ML and biology^4–6^. The breadth of this gap means that biologists often lack the computational skills that are prerequisites for existing ML educational resources^7^. This gap can lead to missed insights from biological data^8^ and contributes to the improper use of ML in biology^1,9^.

Many resources have been created to help researchers acquire skills in ML. Comprehensive resources such as textbooks^10–12^ and online courses require significant time investment, which may not be feasible for active researchers, and teach to a depth that is often unneeded for biological researchers. Other resources such as graphical research and education tools^13–16^, workshops^17^, and written guides and reviews^18–20^, still often focus on coding, mathematics, or running ML. While these resources are important, not all biological researchers will necessarily need to code and run ML experiments independently^21^.

Thus, resources are needed that provide researchers with skills to productively navigate partnerships and collaborations around ML without necessarily directly executing ML research themselves. When designing these resources, it is essential to consider what skills are needed to interpret research that involves ML, communicate with collaborators about ML, and identify biological questions ML can solve. Resources centered around coding, the mathematical underpinnings of ML, or practical advice for using a certain technique do not necessarily fulfill this role.

We created the Machine Learning for Biologists (ML4Bio) workshop to introduce ML to biological researchers. The workshop aims to provide the skills biologists need to be active researchers in a landscape where ML is increasingly prevalent. It focuses on practical research skills such as reading academic papers that use ML and drawing conclusions from ML experiments. We designed the ML4Bio workshop to be approachable and a reasonable time investment; it requires minimal mathematical and computational background and runs for 5 hours over 2 days. A key feature of the ML4Bio workshop is custom software based on the scikit-learn^22^ library, which allows participants to explore and experiment with classification through a graphical interface without computational fluency.

The target audience for the ML4Bio workshop is biological researchers with no computational experience. This audience is somewhat similar to the “discovery biologist” career profile defined by the International Society for Computational Biology (ISCB). However, the workshop emphasizes a more focused set of competencies than all those relevant to this profile^23^.

We used an iterative design process to refine the workshop over a series of 5 sessions from 2018 to 2021. The first 4 sessions were conducted in person, and the fifth was conducted online over Zoom. These iterations gave us insight into how to better align the workshop to our overall goals and address the needs of the workshop’s audience. We then evaluated the effectiveness of the workshop over 3 additional sessions in 2021.

Participants were generally able to achieve the learning goals of the ML4Bio workshop and especially reported an increase in self-reported beliefs that they can engage with ML research. We feel that this success hinges on the workshop’s approachability, careful design, and flexibility. The ML4Bio workshop effectively introduces ML to biological researchers, preparing them for future learning, collaboration, and comprehension of ML experiments in biological domains.

## 2 Workshop Design

### 2.1 Learning Goals

The ML4Bio workshop began with the intention to create a short, intensive workshop that empowers biological researchers to operate in fields where ML is increasingly common and identify where they might pursue ML collaborations in their own research. Rather than tackling the entire field of ML, we chose to focus on classification to limit the scope of the workshop to 1-2 days. The original topics we selected for the workshop involved identifying problems in computational biology, understanding all parts of a typical ML workflow, being able to compare specific classifiers, performing model selection, evaluating a model on new data, and judging the use of ML in biological contexts. These topics were defined based on our professional experiences interacting with biological researchers around ML and through our observations of common challenges in published biological papers that use ML.

Early iterations of the workshop using those topics revealed (a) a mismatch between the selected topics and the coding and mathematical background that is typical of our biological researcher audience and (b) an incorrect scope of the selected topics (too large). To remedy these problems, we employed backward design^24^ to construct realistic learning goals, create assessments for those goals, and develop activities to support participants in achieving those goals. In that process, we focused on preparedness for ML research instead of fully equipping participants to perform ML research independently. The result was the following 4 learning goals whose justification and purpose we discuss in detail below. ML4Bio workshop participants should be able to:

1. Identify machine learning applications and differentiate aspects of a machine learning workflow.
2. Examine a machine learning problem for common factors that influence model selection and problem difficulty.
3. Discover major methodological flaws in a machine learning experiment presented in an academic paper.
4. Demonstrate the belief that they can engage with research that uses machine learning in biology.

#### Learning Goal 1

Characterizing common steps of ML workflows—data pre-processing, training and model selection, and testing and evaluation—gives participants a basis for understanding how ML works and provides a framework for dissecting and understanding unfamiliar ML concepts in the future. Thus, we consider characterizing a ML workflow as an important objective for preparing participants. Additionally, while familiarity with ML terminology is important for research comprehension and communication with collaborators, participants do not need to deeply know all ML terminology by the end of the workshop. As long as participants can generally identify parts of ML and a ML workflow, they are prepared to learn the terminology that is used by their collaborators and is most relevant to their research. This learning goal is somewhat similar to ISCB competency KD3-4^23^, knowledge of “experimental design to ensure the statistical validity of high-throughput experiments.”

#### Learning Goal 2

Specific classifiers are another area of ML that required careful consideration. We originally chose a number of classifiers that we felt were a good introduction to the types of classifiers available and their limitations. General knowledge of what classifiers can and cannot do, and facets of problems such as linear separability that affect model selection, are required to evaluate problem difficulty, but detailed knowledge of specific classifiers is not. Ultimately, we felt that while the classifiers we had chosen do help demonstrate classifiers’ range and limits, participants’ general understanding of the factors that influence model selection will help them irrespective of which classifiers are popular in problems they are interested in. This learning goal is somewhat similar to ISCB competency KC3-5^23^, knowledge of “the suitability of bioinformatics approaches to discovery activities in the life sciences.”

#### Learning Goal 3

ML in biological applications often lacks proper validation or experimental design, especially when those who use it lack a technical background^25,26^. Thus, we consider the ability to find major flaws in a ML experiment, as presented in a research paper, an important part of preparing participants. Since we focus on assessing instead of performing experiments, we teach the types of evidence presented in a paper that indicate overfitting, data leakage, or improper evaluation metrics. However, subtle errors in a ML workflow, such as indirect data leakage, are difficult to find. Researchers whose primary field is ML often miss indirect data leakage, and consistently detecting data leakage is considered an open challenge in ML^26,27^. Therefore, while participants learn the process of assessing a ML workflow, expecting them to be able to consistently find all subtle methodological errors is likely unrealistic. This learning goal is similar to ISCB competency SC3-3^23^, “Critiques experimental design and draws conclusions supported by the data,” and also touches on competency AD3-3, “Is conscious of the risks of overfitting and of appropriate methods for validation and control of overfitting.”

#### Learning Goal 4

Finally, a major focus of the workshop is the affective objective of demonstrating a belief that they can engage in research that uses ML. Affective learning outcomes are those that, as opposed to skills or knowledge, relate to individual dispositions, willingness, preferences, and enjoyment^28,29^. Given that the overarching purpose of the workshop is for biologists to consider pursuing ML as a possible way to solve a problem, including this affective goal is critical to assessing our success. Specifically, while we do not expect them to feel like confident experts, we want participants to believe they can pursue collaborations with ML experts when they identify a problem well-suited to ML. This learning goal is similar to ISCB competency AC3-3^23^, “Demonstrates willingness to incorporate bioinformatics into research.”

The resulting ML4Bio workshop that addresses these 4 learning goals is an online 5-hour workshop divided evenly between 2 days. Data from the initial workshops, coupled with our analysis of the strengths and weaknesses of existing ML resources, led us to follow 3 key design principles^30^ for our workshop. Specifically, we were committed to our workshop (a) focusing on preparedness over fluency or expertise, (b) necessitating minimal coding and mathematical background, and (c) requiring low time investment. The workshop format is a mixture of software activities, active learning^31^ activities, and lectures. Participants use their personal computers to follow along with the online workshop materials and run the ml4bio software. The workshop introduces supervised ML workflows, evaluation metrics, a few common classifiers, and how ML experiments are presented in biological literature. It does not teach participants how to perform ML on their own. Figure 1 shows the various workshop activities and how they relate to the 4 learning goals. Below we explore 3 key features of the workshop design (software, active learning, and drawing on prior knowledge) that support participants in achieving the learning goals.

**Figure 1.**
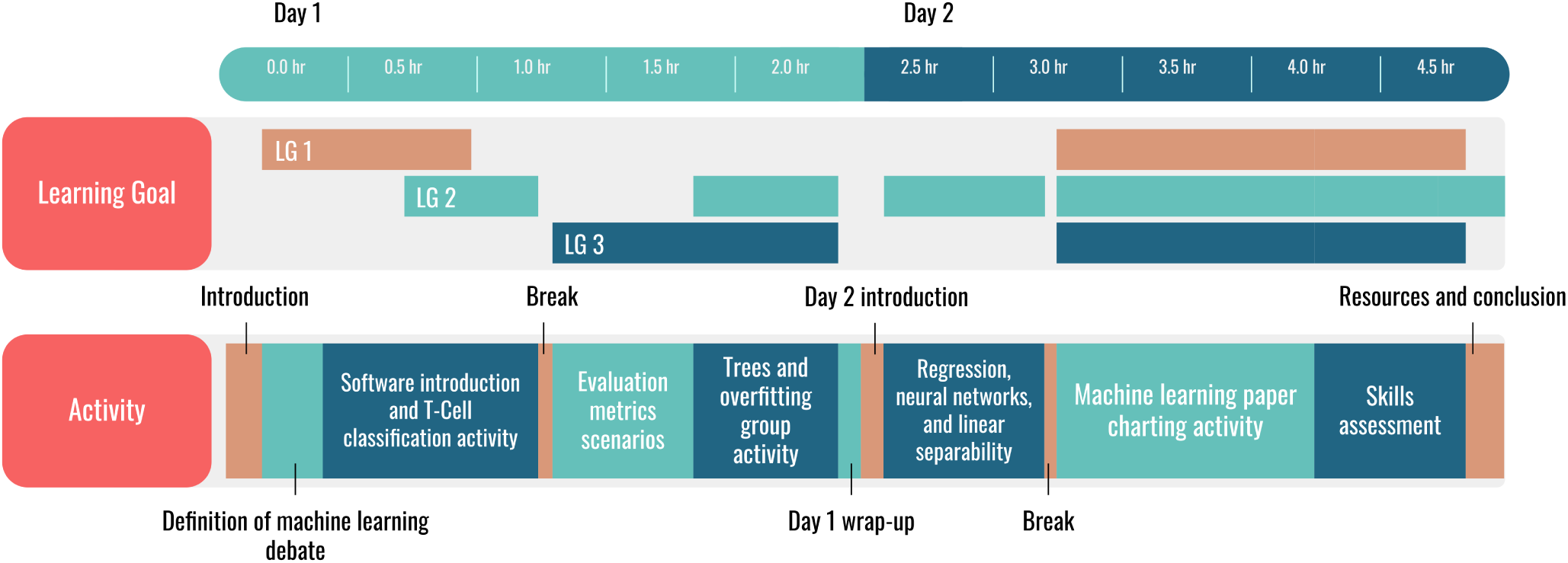
Timeline of the ML4Bio workshop. Activities are shown in addition to which non-affective learning goals (LGs) are addressed by that activity, as defined in Section 2.1.

### 2.2 Software Design

The first—-and perhaps most fundamental—learning goal of the workshop involves understanding a ML workflow. In fact, without understanding that workflow, participants cannot successfully achieve the other objectives. As a result, we wanted to create structural supports in the workshop for this learning goal. Specifically, we wanted to scaffold participants’ learning about the workflow in a way that did not rely on field-specific terminology or existing computational skills. To do so, we created the ml4bio software so biological researchers could visually explore the ML workflow (Figure 2).

**Figure 2.**
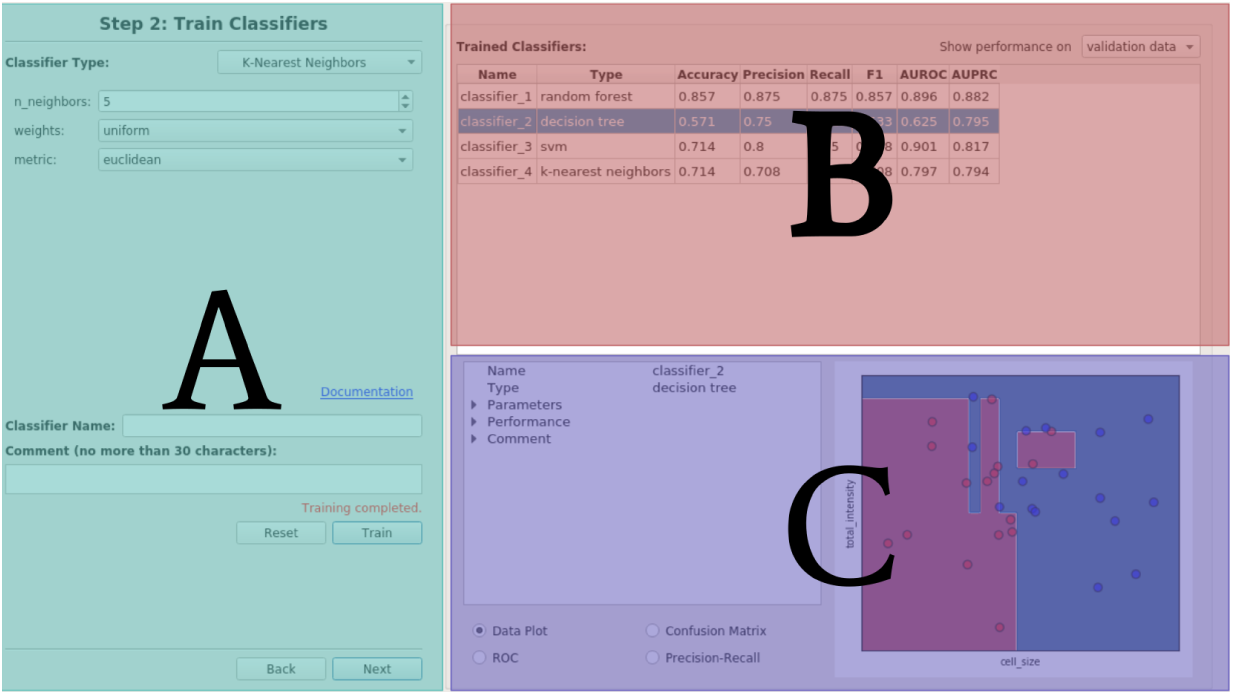
The layout of the ml4bio software interface, with colored panels showing its main sections. Users navigate a ML workflow in panel A, view summarized results in panel B, and view detailed information and data visualizations in panel C.

The ml4bio software is written in Python using the popular ML library scikit-learn^22^. It uses PyQt5 (v5.15.4) for the graphical user interface. Participants are asked to download and install the Anaconda Python distribution and the ml4bio software before the workshop using step-by-step instructions provided on the workshop’s website. We use Anaconda to create a conda Python environment for the ml4bio software via a script that installs and runs the software. In doing so, our software instantiates our design principle of minimizing the need for extensive coding background. The ml4bio package is also available from GitHub (https://github.com/gitter-lab/ml4bio) or PyPI (https://pypi.org/project/ml4bio/).

Once installed, participants and instructors use the software throughout the workshop to walk through ML workflows, compare models and hyperparameters, and visualize decision boundaries and model performance. The software’s user experience is optimized for education instead of other similar software that is designed to perform research-quality data analyses. Workshop participants are warned that the software is not meant to be used in research and is an educational tool. We purposefully limit certain user actions to encourage correct experimental setup and only show a subset of models and hyperparameters to avoid overwhelming users. These restrictions are consistent with our design focus on preparedness (rather than expertise) and low time investment.

The software’s interface is laid out into the left and right halves of the screen (Figure 2). The left half lets the user navigate through the steps of a ML workflow: data selection, training, and testing/predicting, thus directly supporting learning goal #1 (Figure 3). Laying out each of these steps is a key part of the software’s design. At each step, the user is presented with reasonable choices for how to proceed to the next step of the workflow. The software allows users to move forward to the next step, but users generally cannot go back a step without fully resetting and choosing a new dataset. This prevents users from accidentally causing data leakage by performing additional model selection after viewing test set performance or choosing a different test set that might include data from a previous training set. Thus, the user can only perform a complete and standard ML workflow using the software, reinforcing the purpose and flow of each step.

**Figure 3.**
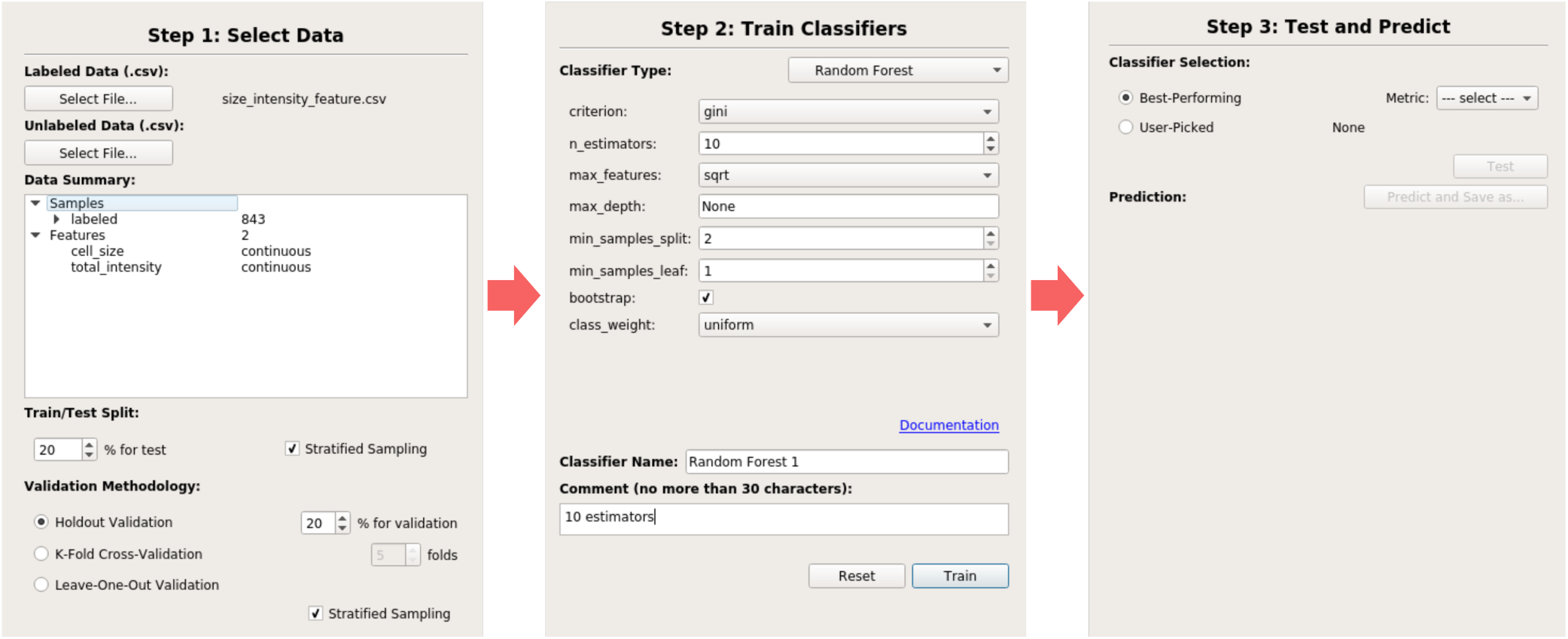
Different configurations of the left half of the software interface throughout a ML workflow.

When <u>selecting data</u>, users can view a summary of the data instances and features in a data summary window. The data is assumed to already be pre-processed. This mirrors our decision to keep detailed data pre-processing methods outside of the scope of the ML4Bio workshop, as pre-processing methods are often domain specific. Users can select a data splitting strategy for both a final test set and a validation set for model selection and whether to use stratified sampling.

In the <u>training</u> step, users can train and compare different classifier and hyperparameter performance on their training set and validation set. The software includes popular classifiers such as decision trees, random forests, support vector machines, neural networks, k-nearest neighbors, and logistic regression. A subset of hyperparameters available in scikit-learn can be changed for each model, and each configuration can be given a name and comment.

As each model is trained, it is added to a table summarizing all trained models’ performances (top half of Figure 4), where a number of classification performance metrics can be viewed for each model on either the validation set or the training set. Each model can be selected, where it is then shown in more detail (bottom half of Figure 4). Here, users can choose to view evaluation curves, a confusion matrix, or a plot of the data with the model’s decision boundary for 2D datasets. Throughout the workshop we especially focus on the decision boundary visualization to show the differences between classifiers, the limits of different classifiers, how certain hyperparameters can affect how a classifier learns. This focus allows us to move away from the specific features of each classifier, which we removed based on feedback from early sessions and refining our learning goals, and instead to focus on how different facets of classifiers and problems affect performance (Learning goal #2).

**Figure 4.**
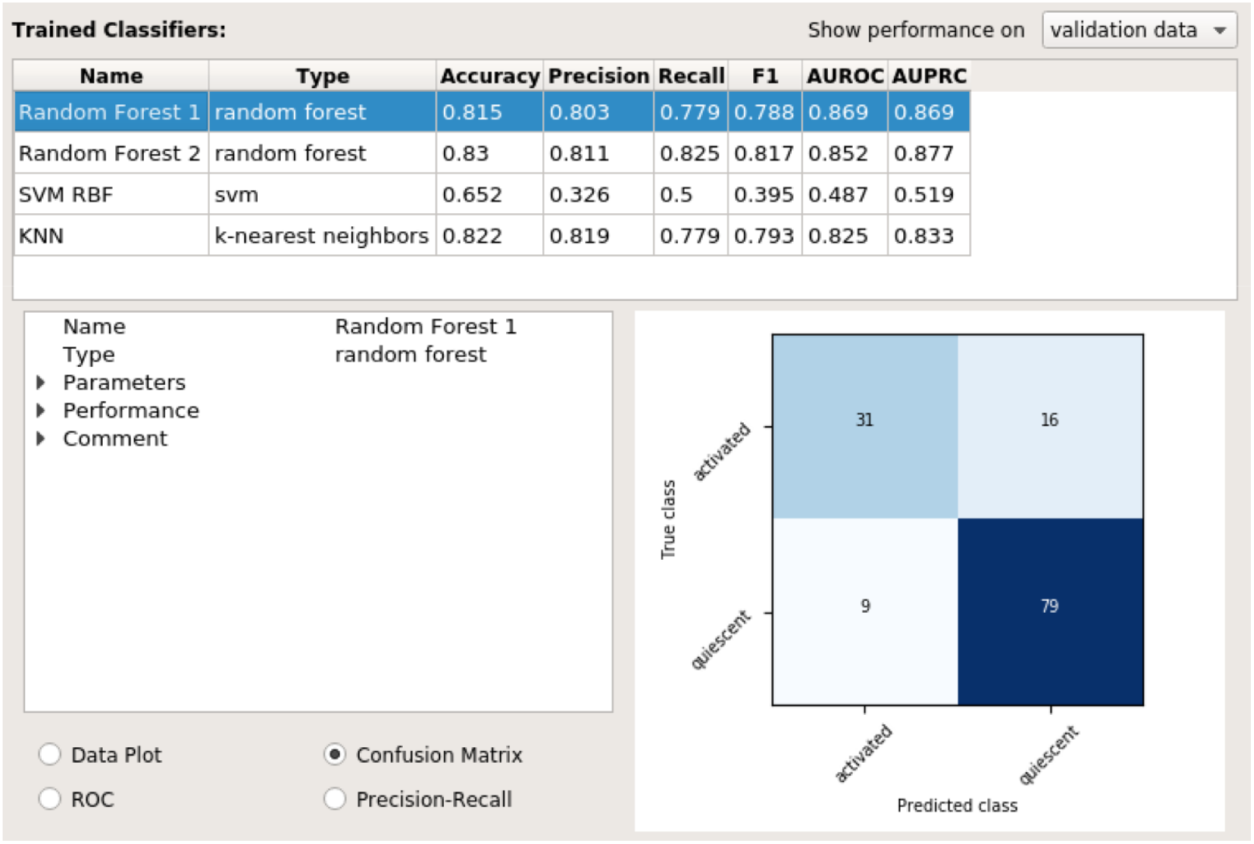
The right half of the ml4bio software interface. The top shows a summary of all classifiers created during model selection, and the bottom shows detailed information on the performance of the selected classifier. Note that multiple classifiers can only be viewed during model selection. The user must select a single model and can no longer see the performance of other models once the test set is examined.

Finally, users can move to <u>testing and predicting</u>. In this step, users can select one of the currently trained models for final classification and evaluation. Users can select a model manually, or select the model that performed best on a certain metric. The software presents a warning that after the test set performance is shown, no more model selection can be performed. After acknowledging this, the right half of the software interface will show only the selected model, but training and validation set performance can still be viewed.

### 2.3 Active Learning

Although our early sessions involved mostly lecture and a summative assessment activity, our use of the backward design paradigm to define learning goals also led us to redesign the workshop activities. Specifically, we transformed the workshop to involve more active learning opportunities for participants^31^.

The workshop now uses a variety of active learning strategies such as scenarios, polls, discussions, and problem solving. It is in an HTML format derived from The Carpentries^32^ lesson template and hosted on GitHub pages (https://carpentries-incubator.github.io/ml4bio-workshop/). This allows anyone to access, propose modifications to, or reuse any workshop materials through GitHub (https://github.com/carpentries-incubator/ml4bio-workshop). Git tags for each workshop date track the evolution of the workshop materials.

As an example of how active learning was added to the workshop, the introductory lesson was changed from a lecture to a debate activity. After sharing a textbook definition of ML^10^, 3 scenarios are presented to participants. One such scenario is a person hand-writing a decision tree from personal knowledge. For each scenario, participants are asked to individually rate how much they do or do not think the scenario is an instance of ML and then justify their position in discussion. We end each scenario by showing how the instructors view the scenario.

We added additional polls and scenarios throughout the workshops. Participants use scenarios and software activities to learn how class balance affects different evaluation metrics. Scenario-based polls are also used as formative assessments at the end of some lessons. These include model selection scenarios, where participants are asked to choose an appropriate classifier based on factors such as the need for interpretability, dataset size, and expected linear separability. In addition, polls at the end of software activities help ensure that participants completed the activity successfully.

Our redesign of the workshop to support participant preparedness (Learning goal #3) led to the largest shift in the workshop towards active learning. Specifically, we added a ML paper charting activity, which is introduced in day 1 and occurs on day 2. On the first day, we ask participants to choose a biological paper that uses ML from a list^33–40^ or a paper they brought, but we make sure that each paper is selected by 2 or more participants. We ask participants to skim this paper before the second day of the workshop.

On the second day of the workshop, after working through an example paper^41^ as a group, participants attempt to fill out a chart cataloging the steps of the ML workflow (Learning goal #1), evaluating model performance (Learning goal #2), and critiquing the experimental design (Learning goal #3). Participants first spend some time individually, then in groups by paper. Finally, we rejoin as a group and discuss issues or interesting conversations that came up.

This activity gives participants guided experience in the interpretation of research involving ML. The transition from learning about ML to having to interpret real ML experiments begins during the workshop. In doing so, we scaffold the participants in moving from lower-levels of Bloom’s taxonomy^42^ to higher-levels, which have been noted to be difficult to teach in ML^43^ in the related Structure of the Observed Learning Outcome taxonomy^44^. Additionally, the choice of paper allows participants to select a paper that interests them. Most of the papers do not cleanly fit into the standard ML workflow taught on the first day, as is expected given the huge variety of ways ML is used. This “messiness” gives participants support in activities that look more like what they will experience in their professional lives.

We found that the online workshop format naturally supported these active learning activities. Zoom polls collected feedback on the scenarios. Screen annotations allowed participants to rate the introductory ML scenarios by drawing on a figure. Breakout rooms facilitated small group discussions of scenarios and the paper charting activity. Participants used the chat to ask questions during lessons that could be answered verbally by the instructor leading the lesson or via chat by a different instructor. Despite these advantages of the online workshops, we observed more extensive and dynamic discussions and research-related questions in our in-person early workshop sessions.

### 2.4 Drawing on Prior Knowledge

Learners come to learning environments with prior knowledge which can both help or hinder their learning^45^. The ML4Bio workshop is no exception: the workshop is intended for those who are involved in biological research, typically graduate students, postdocs, and staff scientists. These participants come to the workshop as trained researchers in a biological domain. Thus, when designing the workshop, we considered an andragogical approach; where *andra*gogy is an approach that specifically focuses on adult learners^46^. Adult learners tend to be motivated by potential applications and learn through drawing on their own prior experiences. We designed workshop lessons to be task-oriented and use real biological applications of ML.

In the second lesson of the workshop, where participants are introduced to the ml4bio software and walk through the ML workflow, we use a motivating example of classifying T-cells as active or quiescent using imaging data^47^. Throughout the workshop, we refer back to this dataset as well as synthetic datasets with the same features and classes that are designed to specifically show some facet of classifier behavior. Other real datasets are included in the ML4Bio GitHub repository from the UCI Machine Learning Repository^48^ and biological studies^49–52^. Using motivating biological problems leverages participants’ prior knowledge to help them understand how classification works. Participants can more easily see what is reasonable or unreasonable in a familiar problem domain. Tailoring ML education to learners’ primary domains has also been effective in undergraduate education^53^.

We use participants’ prior knowledge by centering the academic paper critique activity in the workshop. We expect all participants to be able to interpret and evaluate biological literature. This activity draws on participants’ existing abilities to read and analyze academic papers and merely has them extend those abilities to papers that include ML components. Introducing the skill of reading academic papers from the ground up would take much more than an hour or two to attain^54^. We structured the ML paper charting activity to use this prior knowledge, as participants are encouraged to choose a paper to chart that they are interested in or comes from their research area.

While participants’ prior knowledge generally enhances their learning, we also considered areas where prior knowledge could hinder it. Misconceptions can occur if participants incorrectly apply their prior knowledge and we do not catch and confront the misconception. We were especially cautious when designing the lesson on evaluation metrics. Many of the metrics used to evaluate classifier performance, such as precision and recall, have different meanings in laboratory settings. We directly address this and other possible overlapping terminology to participants.

## 3 Study Design

### Participants

After we reformulated the workshop’s learning goals and activities, we held 3 ML4Bio workshops online over Zoom on May 4^*th*^ and 6^*th*^, August 2^*nd*^ and 5^*th*^, and September 14^*th*^ and 16^*th*^ 2021. Participants were recruited to the workshops via email and could choose to participate in the study after registering for the workshop during the pre-workshop survey. 10, 18, and 19 workshop participants consented to participate in the study in the May, August, and September workshops, respectively. The study was approved by the Institutional Review Board of the University of Wisconsin-Madison (#2021-036), and we obtained electronic informed consent from all study participants. Participants could participate in the workshop and provide informal feedback on the workshop without participating in the study. Here we report only on those who consented to study participation. A breakdown of participant demographics is presented in Table 1.

**Table 1.** Participants of the three workshops. The single “Other” response was noted as “Research Specialist/Technician”. The table includes participants who completed the pre-survey but did not complete the assessment or post-survey.

| Workshop date | Undergraduate | Graduate | Post-doctoral | Staff Scientist | Principal Investigator | Other | Total |
| --- | --- | --- | --- | --- | --- | --- | --- |
| May | 0 | 2 | 5 | 2 | 0 | 1 | 10 |
| August | 2 | 10 | 5 | 0 | 1 | 0 | 18 |
| September | 0 | 9 | 6 | 4 | 0 | 0 | 19 |
| Total | 2 | 21 | 16 | 6 | 1 | 1 | 47 |

### Data Collection

To collect data on the workshop experience and its efficacy in achieving its learning goals, we designed 3 different data collection instruments: a pre-survey, a skills assessment, and a post-survey. The pre-survey was emailed to participants in the week before the workshop; the assessment was given during the workshop; and the post-survey was emailed to participants immediately after the workshop. No directly identifying participant information was collected.

The pre- and post-surveys were designed to collect participant demographic data, record their workshop experiences, and evaluate the workshop’s affective learning goal (#4). In the pre-survey, we collected participant demographic data including current career stage; experience with statistics, ML, coding, and the command line; and overall goals and expectations for the workshop. In the post-survey, participants were asked about their experience in the workshop with regards to their expectations, pacing, time allocation, and general feedback.

The post-survey also included embedded retrospective pre-post designed questions that were used to assess the workshop’s affective learning goal. Retrospective self-assessment has been shown to help prevent response-shift bias^55^, where understanding of the question being asked can change between pre- and post-assessments, while still identifying learning^55^. For instance, a participant’s increased understanding of ML could lead them to realize that they initially understood less than they thought they did, thus resulting in a decrease in self-assessed knowledge after learning. One paired pre-post question was included for verifying the retrospective questions.

In contrast to the surveys, the in-workshop assessment was designed to ascertain whether or not participants had achieved the content learning goals of the workshop (#1-3) and to verify participants’ self-assessment of their knowledge and confidence in ML after the workshop. Participants were given a heavily modified excerpt from a paper that uses random forests to predict microRNA targets^56^. The modifications included feature simplification and the changing model selection to be based on the test set, introducing data leakage into the workflow. Participants were asked to identify parts of the ML experiment such as the model, features, and data splitting strategy and to assess the experiment for overfitting, choice of performance metrics, and data leakage. This assessment allows direct measurement of participants’ ability to understand and assess ML as presented in academic papers. Identifying parts of the ML experiment assesses achievement of learning goal #1, and evaluating the experimental design and model performance assesses achievement of learning goals #2 and #3. Note that without a pre-assessment of learning goals, this assessment strategy does not provide causal evidence that the workshop caused learning goal achievement. We decided that requiring participants to complete a pre-assessment would significantly lower interest in the workshop. Additionally, when paired with the retrospective self-assessment, we can draw conclusions about self-assessment of learning goal achievement and use the in-workshop assessment to verify the level of knowledge post-workshop.

### Data Analysis

The first step in data analysis involved creating matched data sets for each consenting participant. 4 digit codes for each participant linked their pre-survey, in-workshop assessment, and post-survey.

The second step involved analyzing the self-reported survey data. Pre- and post-survey questions related to background, expectations, and experiences (rather than questions related to preparation for future work with ML) were analyzed using basic counts and descriptive statistics. However, both retrospective and paired pre-post questions were compared with two-sided Wilcoxon signed-rank tests. Tests were performed using the scipy.stats.wilcoxon method in SciPy^57^ v1.7.1 with default parameters.

The in-workshop assessment was “graded” for correctness. Workshop designers (who are also ML researchers) determined correct answers for each question on the assessment. Author C.S.M. coded all answers given by participants. Additionally, authors C.S.M. and R.S.R. looked at the participants’ explanations for their responses. From those explanations, we identified common themes in correct (and incorrect) answers. While many questions in the assessment have straightforward answers, later questions are less clear. The final 2 questions in particular, “How well did the model perform?” and “Do you trust the validity of these results?”, do not have an obvious correct answer. We instead compare participants’ responses to possible factors they were asked to identify the presence or absence of in other questions: data leakage, improper performance metrics, and overfitting. How the presence or absence of these factors, and the degree to which they occur, affects participants’ trust in the presented results provides insight into how the participant will engage with ML research.

## 4 Results

### 4.1 Attendees’ Backgrounds and Expectations

Over the 3 workshop sessions there were 47 participants in total who completed the pre-survey. A summary of participants is shown in Table 1. 15 participants only completed the pre-survey, 6 completed only the pre-survey and assessment, and 26 completed all instruments. The 21 incomplete responses include participants who did not return for the second day and participants who completed the workshop but did not fill out the post-survey.

Of the 47 participants, 46 had never taken a ML course, and 6 had never taken a calculus or statistics course. Before the workshop, 13 self-reported as knowing nothing about ML, 27 as knowing a little, and 7 as knowing a moderate amount. Only 3 participants reported having more than a little research experience with ML. Half of participants had at least a moderate amount of coding experience and experience with the command line interface. These data align with our experiences in the initial workshop sessions and provide strong support for our design choice to minimize the need for coding and mathematical background knowledge.

Participants’ expectations generally aligned with the workshop’s learning goals. 33 participants were interested in generally learning about ML with responses such as *“basic overview of machine learning”* and *“understanding how machine learning works*.*”* 24 participants specifically mentioned wanting to learn questions they could answer in their own research using ML or how to apply ML to their research. These expectations align with the current learning goals of the workshop and are consistent with our focus on preparedness rather than ML expertise.

### 4.2 In-Workshop Assessment

In-workshop assessment results (Table 2) show that learning goals #1-#3 were generally achieved, though identifying subtle instances of data leakage proved challenging for many participants. When presented with an altered excerpt from an academic paper, almost all responses correctly identified the target variable, number of instances, model, data splitting strategy, and performance metrics.

**Table 2.**
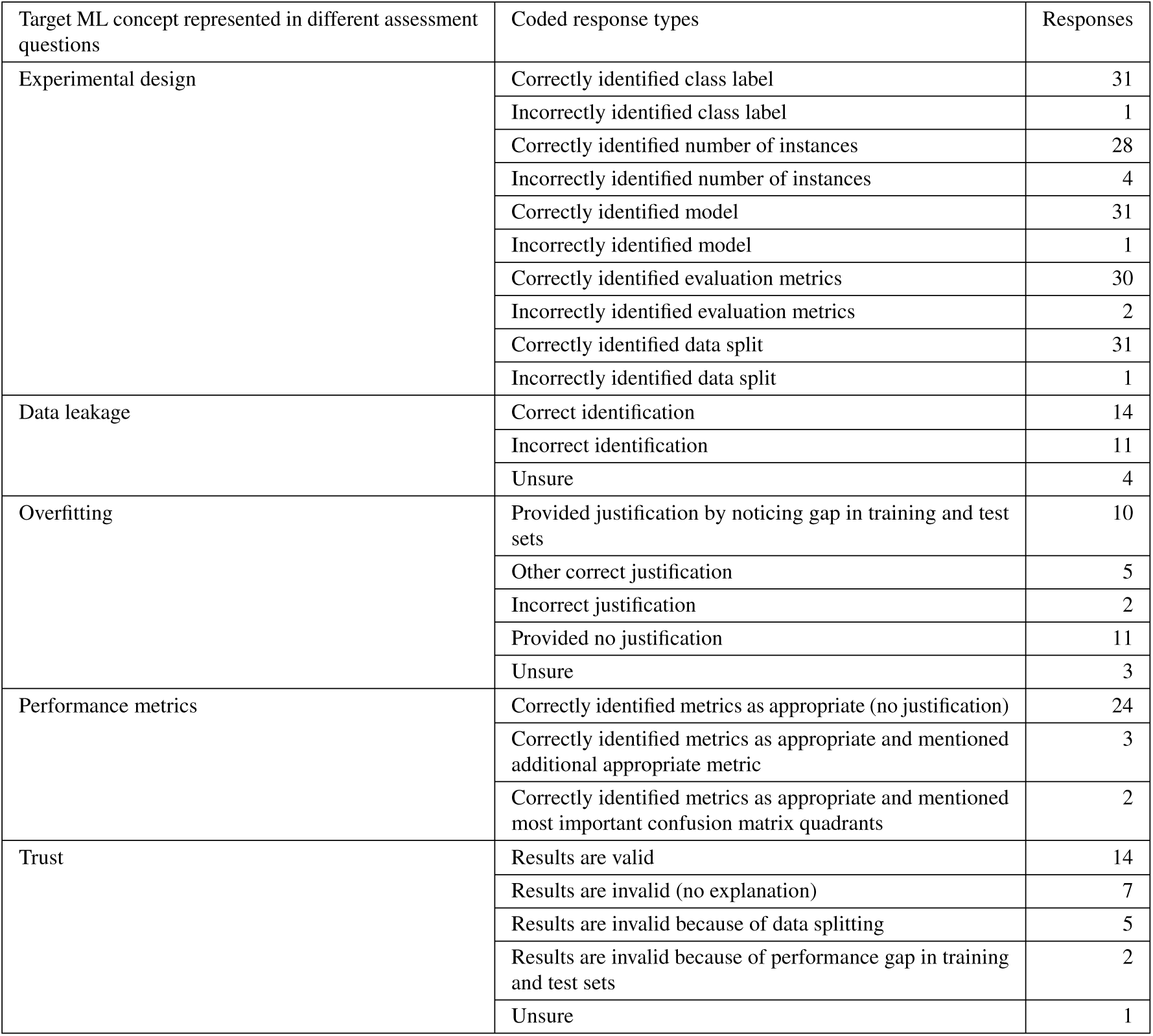
In-workshop assessment results. The numbers presented in this table represent the number of open-ended responses to assessment questions that were coded along each of the dimensions listed. Total number of responses per concept differ because not all respondents answered all of the questions.

| Target ML concept represented in different assessment questions | Coded response types | Responses |
| --- | --- | --- |
| Experimental design | Correctly identified class label | 31 |
|  | Incorrectly identified class label | 1 |
|  | Correctly identified number of instances | 28 |
|  | Incorrectly identified number of instances | 4 |
|  | Correctly identified model | 31 |
|  | Incorrectly identified model | 1 |
|  | Correctly identified evaluation metrics | 30 |
|  | Incorrectly identified evaluation metrics | 2 |
|  | Correctly identified data split | 31 |
|  | Incorrectly identified data split | 1 |
| Data leakage | Correct identification | 14 |
|  | Incorrect identification | 11 |
|  | Unsure | 4 |
| Overfitting | Provided justification by noticing gap in training and test sets | 10 |
|  | Other correct justification | 5 |
|  | Incorrect justification | 2 |
|  | Provided no justification | 11 |
|  | Unsure | 3 |
| Performance metrics | Correctly identified metrics as appropriate (no justification) | 24 |
|  | Correctly identified metrics as appropriate and mentioned additional appropriate metric | 3 |
|  | Correctly identified metrics as appropriate and mentioned most important confusion matrix quadrants | 2 |
| Trust | Results are valid | 14 |
|  | Results are invalid (no explanation) | 7 |
|  | Results are invalid because of data splitting | 5 |
|  | Results are invalid because of performance gap in training and test sets | 2 |
|  | Unsure | 1 |

#### Data Leakage

A methodological error, data leakage, was added to the excerpt. About half of responses correctly identified the presence of data leakage in the experiment (Table 2). Almost all of the responses that provided an explanation for the presence of data leakage correctly cited the lack of validation set or the choice of the final model based on test set performance as evidence of data leakage.

#### Overfitting and Performance Metrics

Most participants who provided an explanation for their response to the presence of overfitting were correct in their reasoning. Participants’ critique of metric choice also showed an understanding of the correct factors to consider, such as how false negatives were more important than false positives in this setting, so sensitivity was an important metric.

#### Trust

Finally, when asked whether or not they trusted the validity of the results, participants had split opinions. Participants who provided an explanation for their response provided correct explanations, such as the data splitting strategy or overfitting as reasons to not trust the results. We are not sure how to interpret other participants’ lack of trust in the paper, and it suggests opportunities for further learning may be necessary to differentiate the severity of different problems with ML workflows.

### 4.3 Affective Outcome

In addition to achieving the content-based learning goals, the data indicates that the workshop was also successful in helping participants achieve the affective learning goal (#4). Recall that our goal here is to support participants in developing the belief that they can engage with ML research. We looked at those beliefs for several domains including training classifiers, reviewing a paper with ML, and identifying a problem well-suited to ML. Based on the self-reported data, participants’ comfort in training classifiers for a research project generally increased after the workshop (*p* = 2.2 × 10^*−*4^, *n* = 26, Wilcoxon signed-rank test) as shown in Figure 5. Before the workshop, over half of participants reported being not at all or a little comfortable, whereas after, among those who responded the majority were either a little or somewhat comfortable. Participants who reported being very comfortable training classifiers after the workshop might show an overestimation of ML skills. We do not expect participants to be able to use ML in their own research without assistance after the workshop.

**Figure 5.**
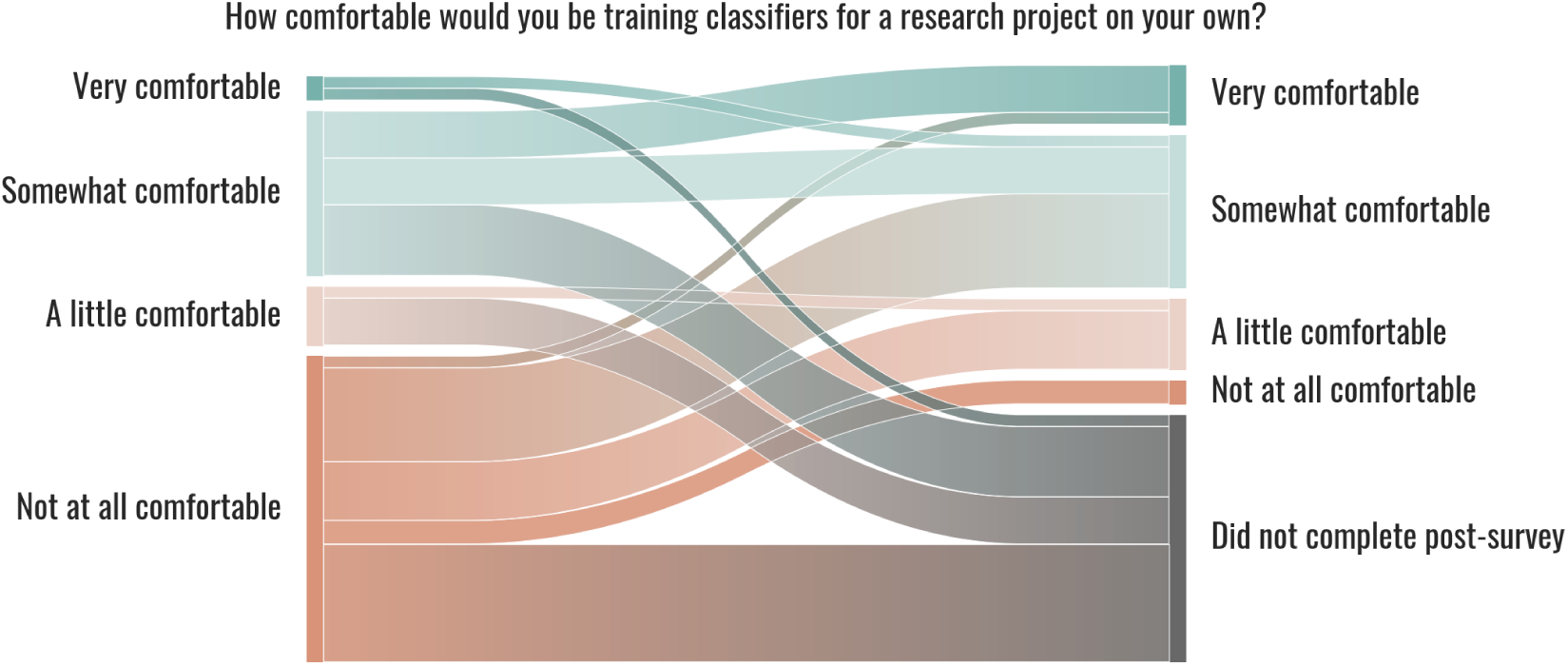
Sankey diagram of participants’ responses pertaining to comfort with ML before and after the workshop across all 3 sessions. Note that a proportion of those who completed a pre-survey and not a post-survey did not attend the workshop at all. 47 completed the pre-survey, 6 completed only the pre-survey and assessment, and 26 completed all 3 instruments.

There was an almost universal increase in self-reported knowledge and confidence from before to after the workshop (Figure 6). We consider these self-reported increases evidence of increased self-efficacy. Participants reported a marked increase in their confidence in identifying a problem that is well-suited to ML in their research (*p* = 5.3 × 10^*−*6^, *n* = 26, Wilcoxon signed-rank test). Participants also reported a significant but lesser increase in general knowledge of ML (*p* = 6.4 × 10^*−*6^, *n* = 26, Wilcoxon signed-rank test) and confidence in reviewing a paper that uses ML (*p* = 1.3 × 10^*−*5^, *n* = 26, Wilcoxon signed-rank test), with the majority of participants reporting that they were somewhat knowledgeable or confident after the workshop. Finally, the most direct evidence we have in participants’ belief that they can engage in ML is the substantial and significant increase in their likelihood to pursue ML for future problems in their work (*p* = 9.4 × 10^*−*6^, *n* = 26, Wilcoxon signed-rank test). A majority of participants reported that they had little or no interest in pursuing and confidence in identifying ML before the workshop, while a majority reported that they were at least very interested and very confident after the workshop.

**Figure 6.**
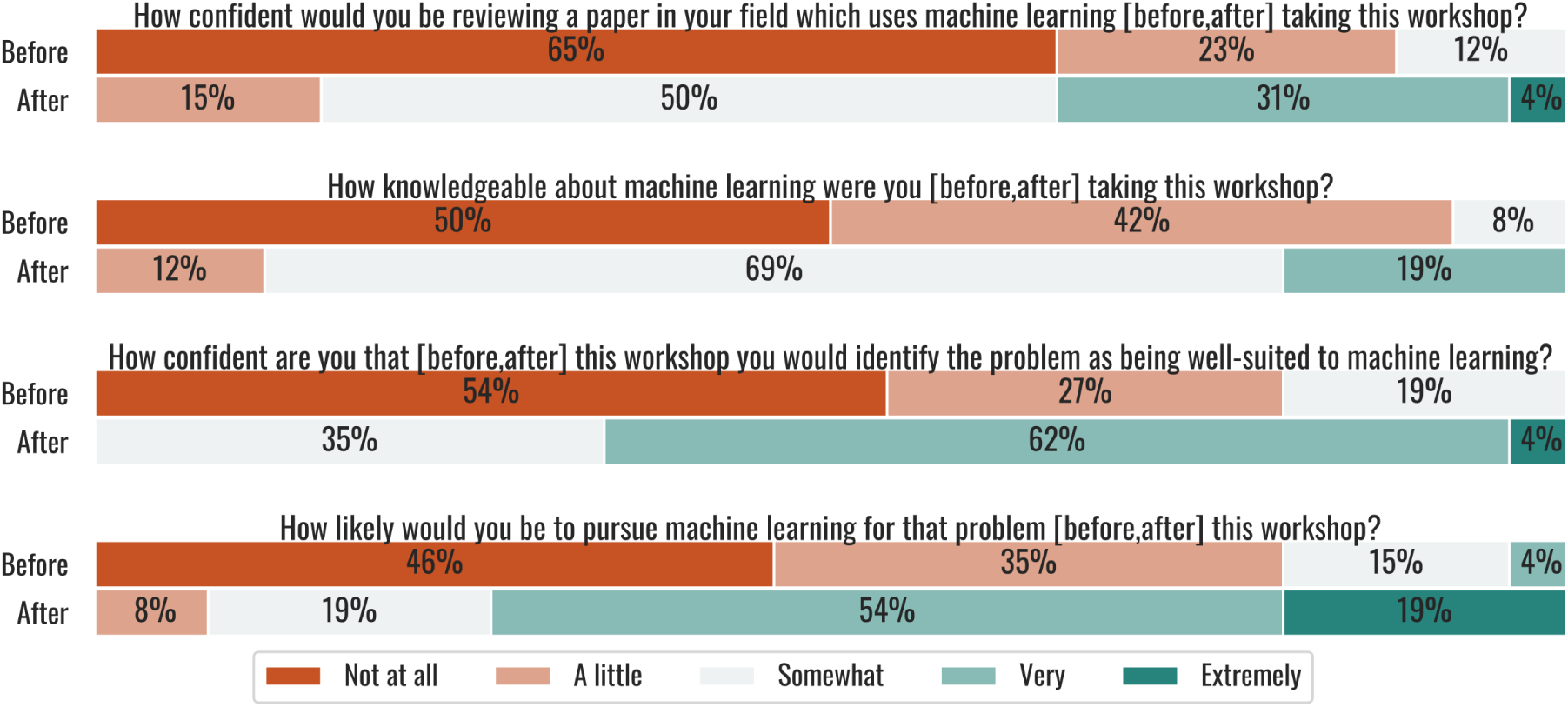
Participant responses to self-reported knowledge, confidence, and interest in ML before and after the workshop. Note that these questions used a retrospective design, meaning that participants were asked about both before and after the workshop in the post-survey.

### 4.4 Workshop Experiences and Expectations

Participants generally reported that the workshop met their expectations. Satisfaction with the workshop was high. 10 participants responded that the workshop exceeded expectations, 10 that the workshop met all expectations, 6 that the workshop met most expectations, and none that the workshop met some or did not meet expectations. There was no clear consensus on which expectations were not met. More real-world examples, coding, clustering, and how to use ML were mentioned.

When asked which workshop topics were most valuable, almost every part of the workshop was named by at least some participants. The two most commonly named parts of the workshop were lessons that involved classifiers and the paper charting activity, with 12 and 7 participants naming those parts, respectively. Participants generally valued learning the variety of classifiers available. Typical responses were focused on *“different classifiers and how they compare”* or *“discussing the different classifiers*.*”* Three participants specifically mentioned that being able to visualize data or how classifiers work was particularly helpful.

A number of the responses mentioning the paper charting activity noted that it was particularly valuable because of its applicability and realism: *“Working through the topics with the ML4bio sample where we could see all the different graphs and assessment statistics to understand how they relate. Also discussing papers where not everything is laid out the same way or fully documented and learning to recognize that*.*” “I thought the research paper exercise was helpful. Specifically because we could bring in articles that interest us*.*”*

Two participants said that the in-workshop assessment was the most valuable part of the workshop for them. Their responses mention that performing an assessment, then checking it as a group, gave immediate feedback on their learning. Other participants noted that scenarios and polls used throughout the workshop also helped them check their own learning: *“I found all of it really valuable. I especially liked the knowledge check at the end through literature evaluations. This course solidified a lot of the ideas behind ML for me*.*”*

Participants were asked which workshop topics they would find difficult to explain at a high level. Details of classifiers were reported as the most common topic participants would have the most trouble explaining. 13 responses mention classifiers in some form in their response. Most responses name specific classifiers and focus on more detail: *“The details of how each algorithm works”, “Logistic regression vs neural networks”*, and *“Explaining how a neural network works*.*”* Other common areas of confusion were model selection and data leakage, with 5 and 3 responses mentioning them, respectively. Responses mentioning model selection often included choosing a classifier for a certain task and optimizing hyperparameters. Despite these continued areas of confusion, participants highlight that the workshop met their goals and provided them with valuable skills and knowledge around ML.

## 5 Discussion

The ML4Bio workshop effectively achieves its goals. Participants left the workshop having attained basic knowledge of ML, increased interest in ML, and preparedness for continued learning. While participants struggled with some of the more subtle aspects of designing ML experiments, namely detecting data leakage, fully preparing participants to perform ML research on their own is not the goal of the workshop. However, improving the workshop such that participants are better able to evaluate ML experiments, and especially detect data leakage, is an important future goal.

### 5.1 Lessons Learned

#### Provide Flexibility for Participants to Bring Their Own Goals

When asked in the pre-survey what they were hoping to learn from the workshop, most participants named a specific research question they were interested in exploring with ML. This aligns with previous literature on adult learners; adult learners are typically more motivated than non-adult learners by real problems^46^. Thus, we found it especially important to ground workshop activities in real or at least realistic data wherever possible.

While having a problem in mind for the workshop can motivate participants, it also complicates meeting participant expectations. The research questions participants bring likely require tools and knowledge beyond a general introduction to ML. Therefore, in workshops geared towards active researchers it is more likely that the workshop’s learning goals may not perfectly align with a participant’s needs.

We mitigate this possible misalignment in a number of ways. We provide a variety of applications throughout the workshop, so that participants are likely to see at least one problem that is similar to their research area. However, we do not cover application-specific pre-processing and feature generation. This is evidenced by multiple participants mentioning that they would want more information on image analysis techniques with ML, even though the first presented dataset is an image classification problem.

Additionally, we provide resources for participants to continue learning about ML. These resources allow participants, even if their goals were not fully met during the workshop, to have an accessible next step for learning about their specific application. We present these resources to participants in the final workshop lesson. The resources include online textbooks, a Jupyter notebook demonstrating a ML workflow, ML-focused code tutorials, and Carpentries workshops for participants interested in learning more technical skills. We plan to continue to grow this list of resources as we are presented with new participant interests.

Finally, during the workshop we give participants space to explore what they find to be interesting. This exploration is clearest during the literature charting activity where participants can choose from a variety of papers to investigate or bring their own. Multiple participants found this activity to be the most valuable part of the workshop. Participants can explore a specific application of ML with support from fellow participants and workshop instructors.

Participants bring their own goals to the workshop. Accommodating these goals can be seen as counter to the backwards-design paradigm, where learning goals are chosen in advance. Incorporating application breadth, resources for participants to continue to learn on their own, and flexibility in the workshop structure are effective tools for creating room for these goals while still conducting a learning goal-driven workshop.

#### Assessments are Worth the Time

Participants found both formative and summative assessments valuable throughout the workshop. Despite assessment being a well-known tool in school-based learning environments^58^, we were initially hesitant to include assessments because we felt they might lower interest from participants. However, multiple participants named the in-workshop assessment the most valuable part of the workshop, though this assessment was originally designed for the study and not directly as a learning tool. Reviewing the assessment afterwards allowed participants to catch misconceptions they otherwise would have taken away from the workshop.

Participants mentioned that other in-workshop assessments, polls, and scenarios throughout each lesson were valuable checks of their knowledge. These quick assessments also allowed the workshop instructors to notice and spend extra time on areas participants were especially confused about.

An additional possible positive effect of the assessments was providing a mastery experience: a challenge that is successfully completed, demonstrating improvement. Mastery experiences lead to increased efficacy and confidence^59^. In the final assessment, almost all participants were able to correctly identify parts of the presented ML experiment. This may have helped demonstrate to participants their new knowledge of ML and helped lead to the marked increases in self-rated confidence and knowledge of ML participants expressed.

#### Set Expectations as Clearly as Possible

Every iteration of the workshop included more information about what participants should expect. However, while the number of participants who noted they had misaligned expectations decreased over subsequent iterations of the workshop, every workshop still had some participants who noted something they were hoping or expecting to learn from the workshop that was not covered.

The nature of the workshop’s audience may have exacerbated this issue. Participants may have come to the workshop having already heard of methods that are popular in their field and expected to learn about them during the workshop. Beyond more general expectations, it may have helped set expectations if we had noted specific, popular concepts that would not be covered in the workshop. For example, we originally noted that the workshop would only present supervised learning and not unsupervised learning. However, some participants had heard of clustering and may not have realized that clustering is a part of unsupervised learning, thus still expecting to learn it. Explicitly stating popular terms and buzzwords can help communicate with participants and manage expectations.

Participants were also uncertain about how deeply they should understand some of the workshop content. This confusion was especially apparent during the lesson on logistic regression and neural networks. Multiple participants expressed that they felt they did not understand all the details of how logistic regression and neural networks work in their post-survey. However, we did not expect participants to understand the mathematics behind these models. Better delimiting what we expected participants to learn about these models and what was out of scope may have reduced this confusion.

### 5.2 Future Directions

We plan to continue to refine and expand the ML4Bio workshop. One area of improvement is to further pare down the knowledge participants need to achieve the workshop’s learning goals. While we have already adapted the scope of some lessons, for instance, by removing the mathematical details of how logistic regression works, it would be productive to systematically approach this refinement. This streamlining could free time to explore more nuanced examples of experimental design flaws like data leakage.

Further clarifying the limitations of the workshop and ML in general would aid participants in choosing their next steps. This could include adding additional cautionary language to the concluding lesson and emphasizing the importance of data cleaning and pre-processing, which often require domain-specific strategies, to help participants leave with a correct understanding of their current skills in ML.

While we are able to determine the achievement of learning goals through the current set of surveys and assessment, some of the questions on the assessment do not encourage useful feedback. Questions on whether or not the results are valid or if the metrics are appropriate do not have a definitive answer, but a large proportion of participants answered them with a one-word “yes” or “no”. Ideally these questions would encourage participants to rephrase their answer or to change the question so that it has a clear correct answer. A large proportion of participants’ answers were not useful for analysis, though they did still appear to provide a valuable learning experience for participants.

A tool to export workflows in the ml4bio software to Jupyter notebooks would provide a powerful link to technical skills. Participants interested in coding could see how workflows they perform in the ml4bio software are expressed as code, giving a smoother transition to coding ML workflows. Currently, we provide a Jupyter notebook with Python code demonstrating an example ML workflow similar to those implemented in the ml4bio software. Participants can run this notebook in a web browser with Binder^60^.

A few participants in each workshop consistently struggled with software installation before and during the workshop. Some issues arose from installing and configuring Anaconda. Others were due to the scripts we provided to create the required conda environment or the environment itself. Including more details about expected behavior and installation screenshots in our setup instructions partially alleviated but did not eliminate these issues. A cross-platform installer that provides the ml4bio software and required datasets, possibly using the conda constructor tool, would make the workshop more accessible and reduce the amount of command line troubleshooting required. Rewriting the ml4bio software to run in a web browser would minimize the technical requirements and help scale the workshop to larger audiences. However, this would require substantial software development. There are also numerous ways to improve the ml4bio software such as support for more classifiers, visualization of datasets with more than 2 features, saving models and settings, more interface tooltips, and better text scaling.

We plan to expand our current instructor notes to the point that we could provide the ML4Bio workshop as a full lesson plan others could teach. While all activities in the workshop are laid out in the online materials and current instructor notes, they are not detailed enough for someone to teach the workshop without first observing it. The workshop could also be scaled up to larger sessions if additional helpers were present to lead group activities and troubleshoot software issues. The ML4Bio workshop has joined The Carpentries Incubator to gain additional support and feedback and to expand the audience and possible future instructors. Our workshop design and instruction have already benefited from the principles taught in The Carpentries Instructor Training and specific suggestions from Carpentries instructors. We hope to continue to expand and improve the ML4Bio workshop so that it continues to be an effective tool for helping biologists participate in an increasingly computational research world.

## Acknowledgements

We thank the ML4Bio workshop and study participants; Devin Wixon for early feedback and design help; Chris Endemann, Beth Meyerand, Lauren Michael, Ross Kleiman, Sarah Stevens, and Zijie J Wang for workshop content suggestions; and Karen Zoladz for survey advice. This work was supported by NIH award T15LM007359, NSF award DBI 1553206, the Morgridge Institute for Research, and the University of Wisconsin–Madison Office of the Vice Chancellor for Research and Graduate Education with funding from the Wisconsin Alumni Research Foundation.

